# Crystal structure of a constitutive active mutant of adenosine A_2A_ receptor

**DOI:** 10.1101/2021.09.08.459393

**Authors:** Min Cui, Qingtong Zhou, Yuan Weng, Deqiang Yao, Suwen Zhao, Gaojie Song

**Author notes:** These authors contributed equally to this work (Min Cui, Qingtong Zhou). Corresponding author. (Q.Z.), (G.S.).

## Abstract

Previously we reported a common activation pathway of the class A G protein-coupled receptors (GPCRs) in which a series of conserved residues/motifs undergo conformational change during extracellular agonist binding and finally induce the coupling of intracellular G protein, and successfully predicted several novel constitutive active or inactive mutations for A_2A_ adenosine receptor (A_2A_AR) through this mechanism (Zhou et al., 2019). Here we determined the crystal structure of a typical A_2A_AR constitutive active mutant I92N in complex with agonist UK-432097 to reveal the molecular mechanism of mutation-induced constitutive activity. The mutated I92N forms a hydrophilic interaction network with nearby residues including W^6.48^ of the CWxP motif, which is absent in the wild type A_2A_AR. Although the mutant structure is overall similar to the previously determined intermediate state A_2A_AR structure (PDB ID: 3QAK), the molecular dynamics simulations suggested that the I92N mutant stabilizes the metastable intermediate state through the hydrophilic interaction network and favors the receptor conformational transition towards the active state. This research provides a structural template toward the special pharmacological outcome triggered by conformational mutation and sheds light on future structural or pharmacological studies among class A GPCRs.

## Introduction

GPCRs are a group of seven-transmembrane (7TM) proteins that can sense extracellular chemical/light/odor signals to transduce downstream cellular adaptors such as G proteins (Venkatakrishnan et al., 2013; Yang et al., 2021). The signal transduction is accomplished through binding of its agonist in the extracellular pocket that triggers conformational changes of the 7TM, which in turn create enough space in the intercellular region to accommodate G protein (Rasmussen et al., 2011; Weis & Kobilka, 2018). While the ligand-receptor binding modes are varied among different receptors, the transition pathways from a ligand-free, inactive state to both agonist- and G protein-bound, active state are roughly similar in the intercellular side, and characterized by a narrow inward movement of TM7 toward TM3 and a wide outward movement of TM6 in their cytoplasmic ends. In addition to the active and the inactive snapshots determined by crystallography or cryo-EM, there are also a series of intermediate states during the conformational transition, which has also been well illustrated by NMR studies (Manglik et al., 2015; Ye, Van Eps, Zimmer, Ernst, & Prosser, 2016). Of note, the special role of the sodium-coordinating residues (such as D^2.50^ and S^3.39^)(Ballesteros–Weinstein numbering in superscript) on receptor stability and G protein signaling have been comprehensively explored from the aspects of structural biology (Ballesteros & Weinstein, 1995; White et al., 2018) and biophysics (Eddy et al., 2018; Lee, Nivedha, Tate, & Vaidehi, 2019; Song, Yen, Robinson, & Sansom, 2019), while the importance of other motif residues (such as CWxP, PIF and DRY) are yet to be completed. We have previously reported a common GPCR activation pathway that directly links the ligand-binding pocket with the G protein-binding region (Zhou et al., 2019), this common activation mechanism is featured by switching or repacking of dozens of paired residues within the intercellular half of the 7TM including those conserved class A motifs (Erlandson, McMahon, & Kruse, 2018; Thal, Glukhova, Sexton, & Christopoulos, 2018). This mechanism is confirmed by designing of constitutive active or inactive mutations within the pathway. The adenosine A_2A_ receptor (A_2A_AR) is a prototypical member of the class A subfamily of the G protein-coupled receptors (GPCRs) that is widely distributed in various tissues and organs of the human body and participating in many important signal regulation processes. Using A_2A_AR as an example, we have functionally validated 6 mutations with increased basal activity (*i*.*e*., constitutive active mutations) and 15 mutations with decreased or abolished activity (*i*.*e*., constitutive active mutations)(Zhou et al., 2019).

Abnormal function of A_2A_AR has been linked to neurodegenerative diseases such as Parkinson’s disease, Huntington’s disease, inflammation and coronary heart disease. A_2A_AR has been considered as a prototypical receptor in the GPCR structural biology field and dozens of A_2A_AR structures with different types of ligands and/or adaptors have been determined. A_2A_AR in complex with antagonist was crystallized to inactive state (Liu et al., 2012), while these agonist-bound A_2A_ARs were mostly crystallized to the intermediate state (Lebon, Edwards, Leslie, & Tate, 2015; Lebon et al., 2011; F. Xu et al., 2011). Compared to the full active conformation that acquired with the presence of G protein or mini-G protein (Carpenter, Nehme, Warne, Leslie, & Tate, 2016; Garcia-Nafria, Lee, Bai, Carpenter, & Tate, 2018), the intermediate state is within the receptor’s transition pathway from inactive state to active state. To investigate whether these constitutive active mutations are linked to unobserved conformational states of A_2A_AR, we tried to crystallize agonist bound A_2A_AR in combination with different constitutive active mutations (Zhou et al., 2019): I92^3.40^N, L95^3.43^A and I238^6.40^Y Among them, the I92^3.40^N was predicted to form a hydrogen bond with W246^6.48^, while L95^3.43^A and I238^6.40^Y were thought to loosen the hydrophobic lock between L95^3.43^, I238^6.40^ and V239^6.41^. All these mutations are hypothesized to favor the active conformation by rotating the intercellular half of TM6, which thus loosen the TM3–TM6 contacts to allow TM6 moving outward in an easier way to create enough space for recruiting of downstream G protein.

## Results

All mutations were made based on previous crystallized A_2A_AR construct with the third intracellular loop (ICL3) replaced with BRIL (Liu et al., 2012)(referred as wild-type A_2A_AR hereafter unless further mentioned). These variants were expressed in insect cells and purified to similar homogeneity as wild-type (WT) A_2A_AR (Figure 1A-B). We firstly measured their thermal stabilities in *apo* state or in complex with agonist (CGS21680) or antagonist (ZM241385) by CPM-based thermal-shift assay (Figure 1C). All *apo* variants were relatively unstable without the presence of ligands. Interestingly, the WT A_2A_AR showed the best thermal stabilities in all conditions compared to these variants. Especially, the I92^3.40^N and L95^3.43^A *apo* proteins each showed a significantly decreased melting temperature (3∼5°C) compared to the WT, whereas the decreases can be fully retrieved with presence of CGS21680 but only partially retrieved by ZM241385 (Figure 1D). The result suggested that these mutations indeed break the equilibrium formed by the WT A_2A_AR and driven the receptor from inactive to active state, this can also be testified by the fact that these mutants increased the basal activities by 7-28 fold (Zhou et al., 2019). These *apo* variants are metastable but can be stabilized by the agonists who favor the active state, whereas the antagonists who lean toward the inactive state are incompatible with these mutations. Compared to the WT A_2A_AR, the I238^6.40^Y showed comparable melting temperatures in *apo* state or with agonist, but with a decreased melting temperature with antagonist (Figure 1D). Such result suggested that Y238^6.40^ might additionally stabilize the local environment by its bulky aromatic side-chain, a principle that previously characterized and summarized on a class B GPCR, glucagon-like peptide-1 receptor (Y. Xu et al., 2019).

**Figure 1.**
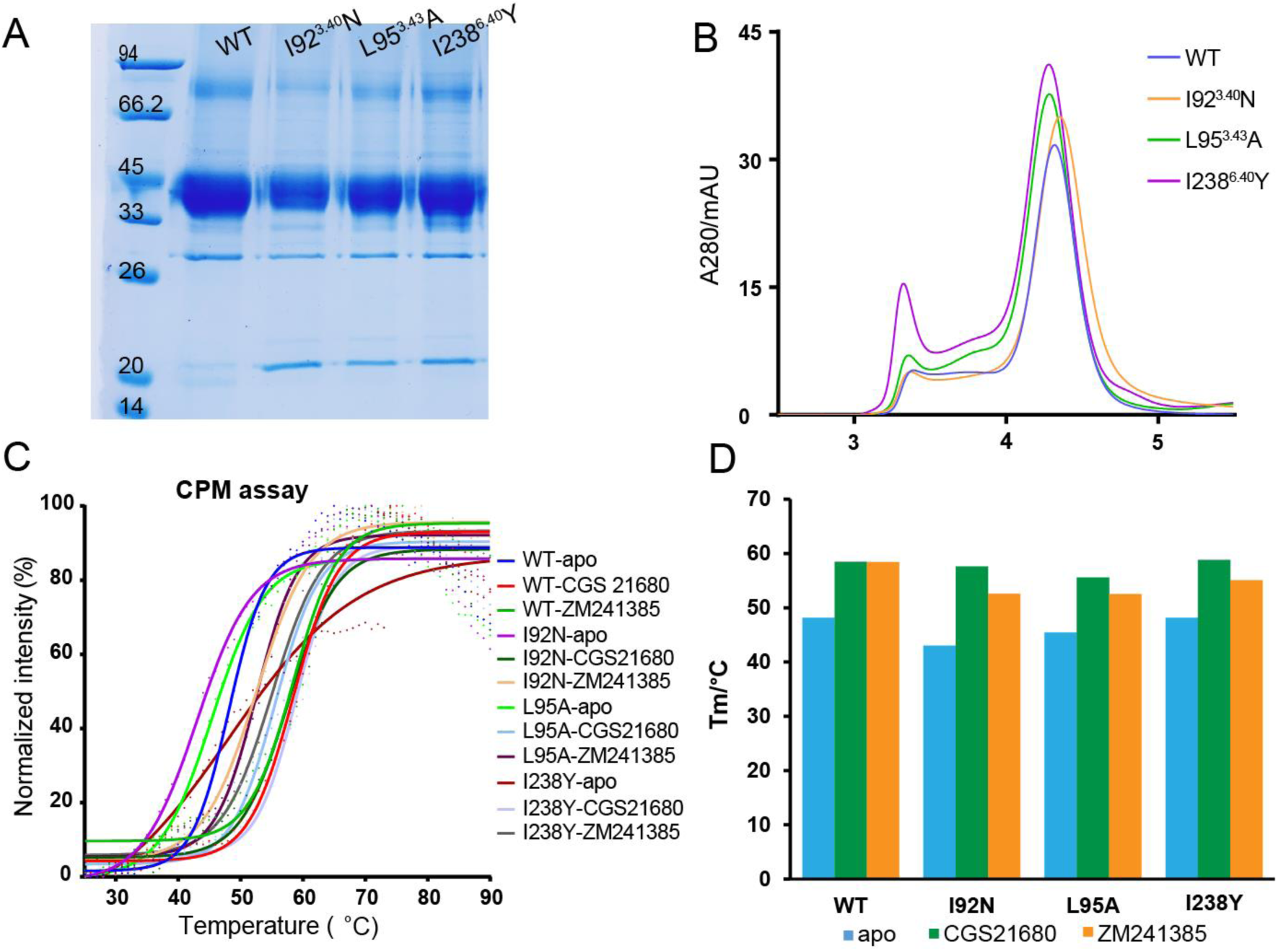
Purification and thermal-shift assay of A_2A_AR constitutive active mutants. (**A**), SDS-PAGE of purified WT and mutant A_2A_AR fusion proteins. (**B**), size-exclusion chromatography (SEC) suggest that the A_2A_AR fusion proteins are mostly monomeric and of similar homogeneity; (**C-D**), Thermal-shift profiles (**C**) and Tm plots (**D**) of A_2A_AR WT and mutants in *apo* state or in complex with agonist (CGS21680) or antagonist (ZM241385). In the thermal-shift assay, 500 mM NaCl was added in parallel to each experimental buffer for strict comparison since sodium is an allosteric effector for A_2A_AR. **Source data 1**. Raw representative western blot of A_2A_AR WT and mutants during purification. **Source data 2**. Raw size-exclusion chromatography data of A_2A_AR WT and mutants. **Source data 3**. Raw CPM-based thermal-shift assay data of A_2A_AR WT and mutants. **Figure supplement 1**. Purification and crystallization of I92N mutant with UK-432097.

Since the agonist performs better than antagonist in thermal-shift assay (and also logically as described), we tried co-crystallization of all three variants with agonist CGS21680 but failed. We then tried co-crystallization with another agonist, UK-432097, who has similar potency as CGS21680 and was the first agonist that crystallized with WT A_2A_AR (F. Xu et al., 2011). We successfully crystallized mutants I92^3.40^N and L95^3.43^A with UK-432097, but can only optimize the crystals of I92^3.40^N to suitable size and collected the data to 3.8 Å (Figure 1—figure supplement 1, Table 1). The structure was determined using previous intermediate state A_2A_AR structure (PDB ID: 3QAK) as searching model.

**Table 1.**
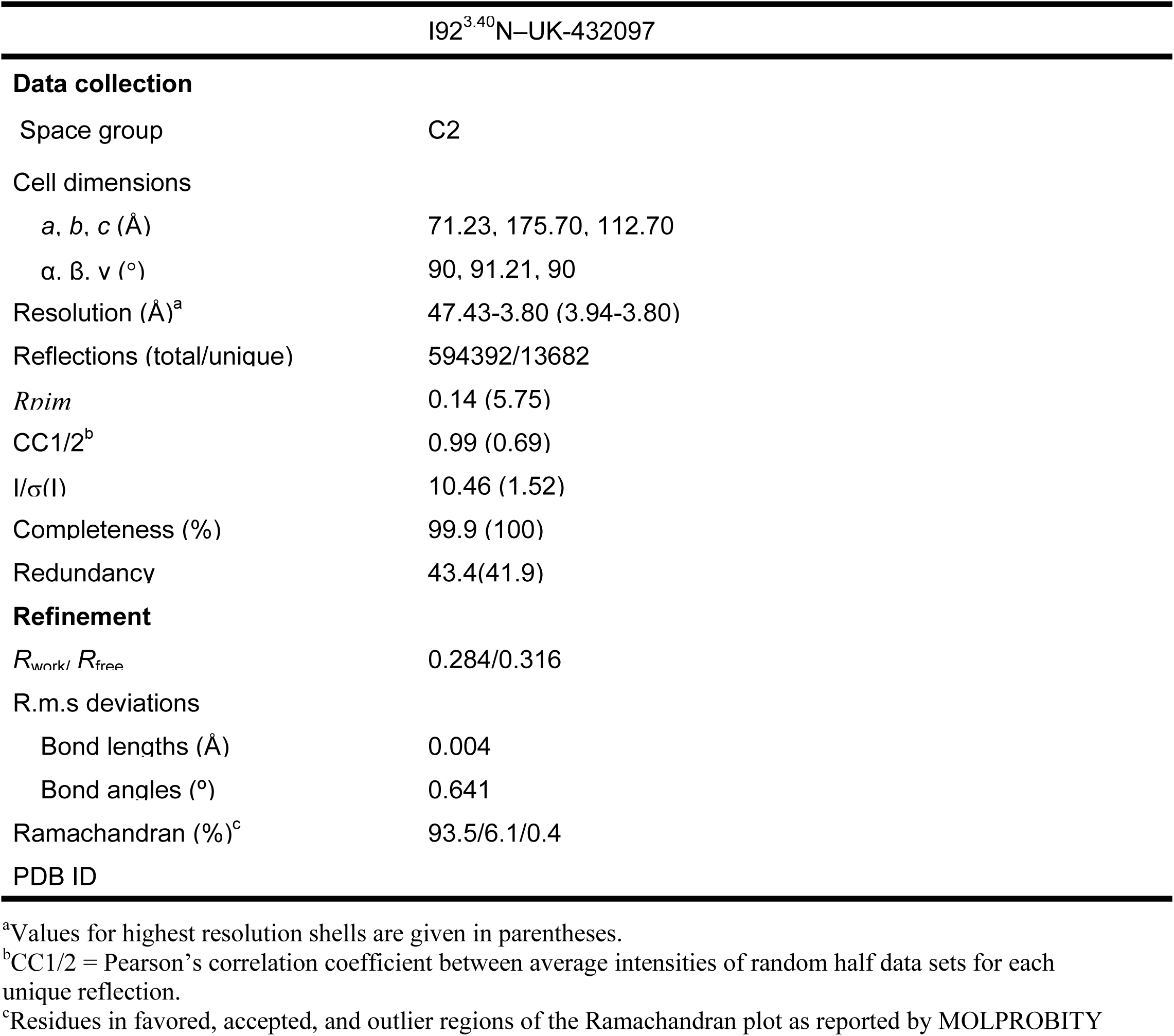
Data collection and refinement statistics.

The I92^3.40^N–UK-432097 structure is overall similar to previous intermediate A_2A_AR structure (F. Xu et al., 2011) with a Cα r.m.s.d of 0.46 Å, and is distinct from the inactive (Liu et al., 2012) (Cα r.m.s.d of 1.74 Å) or active (Carpenter et al., 2016) (Cα r.m.s.d of 1.61 Å) state structure. Consistent with our prediction, in the variant structure the N92^3.40^ forms a hydrogen bond with W246^6.48^ (Figure 2A-B). Unexpectedly, the side-chain of N92^3.40^ also forms a weak hydrogen bond with the carbonyl group of C185^5.46^, as well as an even weaker interaction with the side-chain of N280^7.45^. Of note, all these residues are relatively far away from each other in the inactive state (Erlandson et al., 2018), thus these residues and their local structures undergo conformational change and move together during receptor activation. Obviously, above hydrophilic interactions are not possible to happen in the WT A_2A_AR since the corresponding position of 3.40 is a hydrophobic isoleucine residue (Figure 2C). Actually, I92^3.40^, well-known as part of the P^5.50^-I^3.40^-F^6.44^ motif that triggered signaling initiation(Schonegge et al., 2017; Wacker et al., 2017; Zhou et al., 2019), locates in an edge between the transmission switch and the hydrophobic lock in the structure, whereas in the inactive conformation it involves more with the hydrophobic lock. Therefor it is our estimation that the I92^3.40^N mutation disturbed the local environment in the a*po* (inactive) state, while this disharmony may be compromised through adding the agonist by which induces departure of the N92^3.40^ from the hydrophobic lock and formation of these hydrophilic interactions, these analysis are in line with the thermal-shift assay (Figure 1C,D).

**Figure 2.**
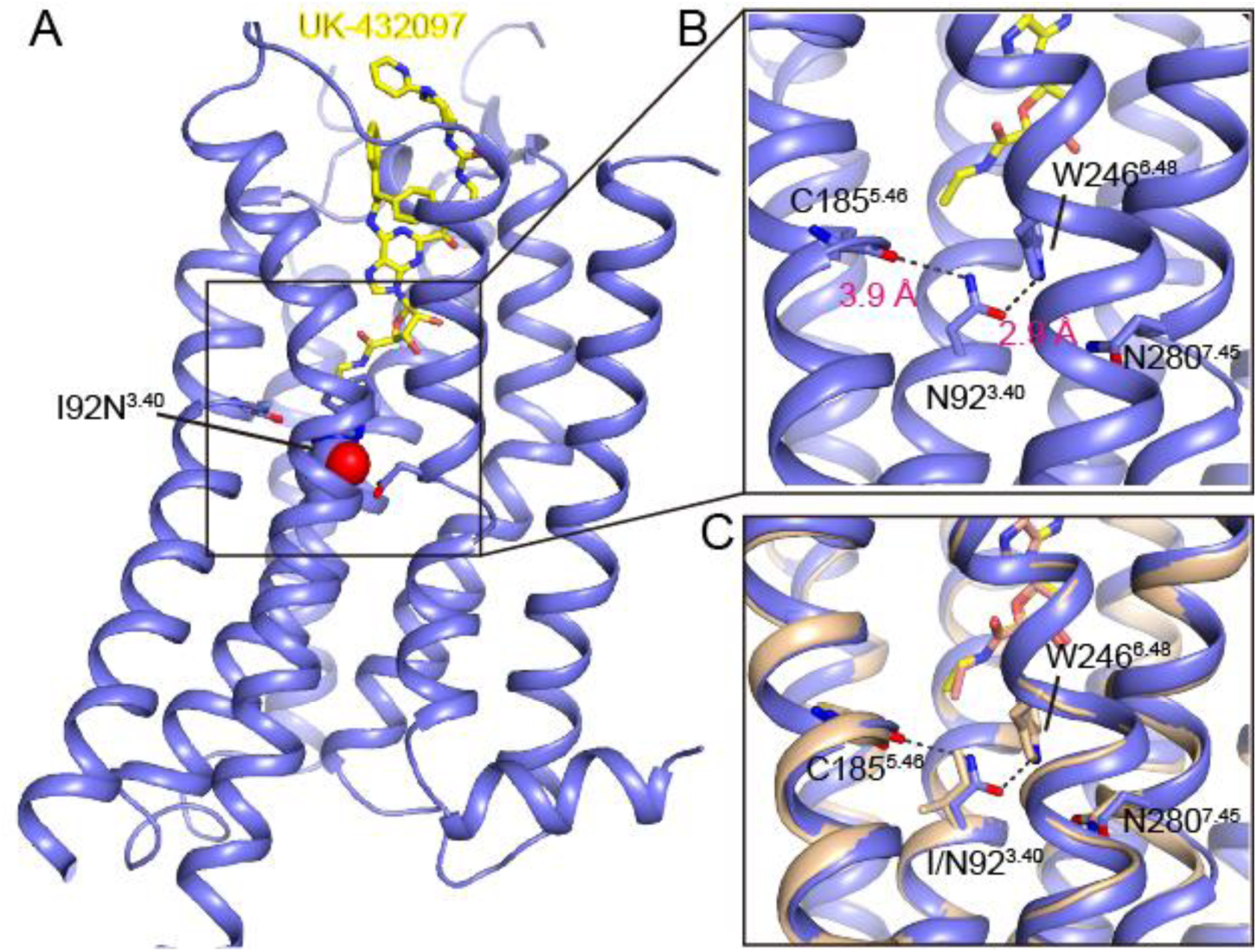
The I92^3.40^N mutant structure of A_2A_AR in complex with UK-432097. (**A**), Overall structure with agonist UK-432097 shown as yellow sticks and N92^3.40^ shown as spheres. (**B**), Zoomed-in view of the region around N92^3.40^ within the mutant structure. Hydrogen bonded residues are labeled and their distances are marked. (**C**), Superposition of the mutant structure with the intermediate state WT structure (PDB ID: 3QAK). In the WT structure, I92^3.40^ and its nearby residues are shown as orange sticks. **Figure supplement 1**. Crystal packing of I92N–UK-432097.

For A_2A_AR, the active state is roughly identical with the intermediate state in the transmission switch region but distinct in the intercellular end of TM6, which moves further outward by >10 Å to accommodate G protein. The transition from intermediate to active state in the intracellular region requires the switch and new interactions formed by key residues R102^3.50^, Y288^7.53^, as well as the residues in G protein. However, in the I92^3.40^N–UK-432097 structure we did not see further conformational change in the intracellular end of TM6 compared to previous intermediate A_2A_AR structure. This is consistent with previous finding that full activation of a GPCR requires engagement of its downstream G protein, as validated in many receptors including A_2A_AR and β_2_AR (Eddy et al., 2018; Nygaard et al., 2013; Thal et al., 2018; Ye et al., 2016).

Crystal structure represents typically a single conformation of individual protein, while it is known that GPCRs are very dynamic and multiple conformations are employed during their physiological events (Latorraca, Venkatakrishnan, & Dror, 2017). To further explore the dynamic events of the variant and previous WT A_2A_AR intermediate structure, we performed all atom molecule dynamics (MD) simulations to monitor how the I92^3.40^N mutation might affect the dynamic or conformation of A_2A_AR (Figure 3-4). All simulations including the I92^3.40^N and WT together with UK-432097 or without UK-432097 (*apo*) were conducted at 1 μs timescale. Structural comparison among the inactive, intermediate and active structures of A_2A_AR reveals the step-wise conformational change occurred in the residues centered at I92^3.40^ (Figure 3A) to final dense packing upon receptor activation, as seem from the decreasing inter-residue minimum distances (Figure 3C). When bound to UK-432097 (trajectory I92N–UK-432097), the mutated N92^3.40^ was mostly stabilized in its original position that identical with the active/intermediate states but distinct from the inactive state (Figure 3A). Quantitatively, the N92^3.40^ reserves its hydrogen bond interactions with C185^5.46^ and W246^6.48^ to percentages of 98% and 78%, respectively (Figure 3B). In the mutant simulation without UK-432097 (trajectory I92N–*apo*), the N92^3.40^–C185^5.46^ interaction is largely disrupted; in contrast, the N92^3.40^–W246^6.48^ interaction is well maintained at early stage and fluctuation happens only at the 2^nd^ half timescale (Figure 3C). For the simulations of WT A_2A_AR, the minimum distances between I92^3.40^ and W246^6.48^/C185^5.46^/N280^7.45^ are also measured. The minimum distances of I92^3.40^–W246^6.48^ and I92^3.40^–C185^5.46^ are roughly stable during the simulation with UK-432097 (trajectory WT–UK-432097); in contrast, with removal of the agonist (trajectory WT–*apo*) both distances are fluctuated and apparently larger than the ones with the presence of UK-432097 (Figure 3C). The I/N92^3.40^–N280^7.45^ distance is overall not that sensitive compared to the other two pairs of distances. However, we can still see that the minimum I92^3.40^–N280^7.45^ distance is apparently larger in average in the simulation of WT–*apo* (without UK-432097) compared to WT–UK-432097, while the N92^3.40^–N280^7.45^ distance is not differentiable between the simulations of I92N–UK-432097 and I92N–*apo* (Figure 3C). All these results indicated that in addition to the agonist who drives the transition of receptor from inactive to intermediate/active state by forming multiple interactions with the pocket residues, the I92^3.40^N also plays an essential role by disturbing the local environment and forming these hydrophilic linkages which escorted conformational change of the intracellular G protein-binding region.

**Figure 3.**
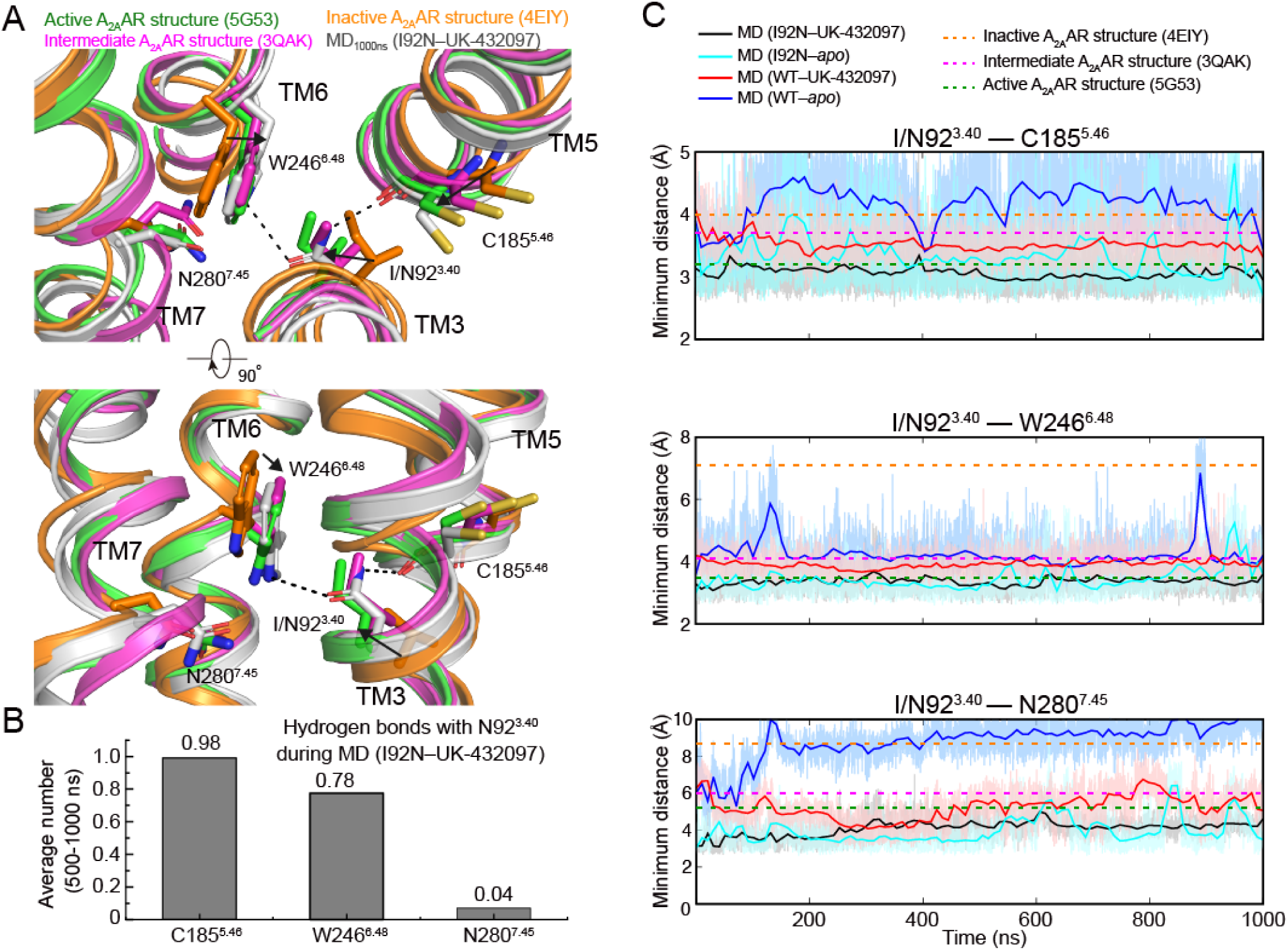
Conformational dynamics of the ligand-binding pocket in the molecular dynamics (MD) simulations of wild-type A_2A_AR and its mutant I92N. (**A**), Structural comparison of the residues around I/N92^3.40^ between MD final snapshot and the released structures of A_2A_AR in different states. Inactive (antagonist-bound), intermediate (agonist-bound) and active (both agonist and G protein bound) A_2A_AR structures are colored in orange, magenta and green, respectively (Carpenter et al., 2016; Liu et al., 2012; F. Xu et al., 2011). (**B**), Statistics of hydrogen bonds between N92 and its surrounding residues during the last 500 ns MD simulation of UK-432097-bound A_2A_AR mutant I92N. (**C**), Representative distances between I/N92^3.40^ and its surrounding residues C185^5.46^, W246^6.48^, and N280^7.45^. Minimum distances were measured between non-hydrogen atoms. Dashed horizontal lines indicate values for the released structure of A_2A_AR in different states (inactive state, orange; intermediate state, magenta; active, green).

At the intracellular region, the distinct performances of conformational dynamics between the WT and mutant receptors during MD simulations suggested unique role played by I/N92^3.40^. Within all four types of trajectories the minimum distances between the ionic lock residues (R102^3.50^ and E228^6.30^) are far less than that of active state (18.8 Å). Nevertheless, while all other trajectories are fluctuating between the inactive and intermediate states, the WT–*apo* apparently returned back to the inactive state after 200 ns of simulation, judging from the steadily formed ionic lock as well as the Cα–Cα distance between R102^3.50^ and the first residue of TM6 (T224^6.26^). Consistently, the solvent-accessible surface area (SASA) of G protein-binding site for the WT–*apo* snapshots are averagely smaller than the other three. The trajectories of I92N–UK-432097 and I92N–*apo* are roughly similar, with far less distances of key residue pairs (R102^3.50^–T224^6.26^, R102^3.50^–E228^6.30^) (Figure 4B, top and middle) but much closer SASA of G protein-binding sites (Figure 4B, bottom) compared to those of active structure. Such asynchronous events between the creation of intracellular cleft for G protein entering and further outward movement of the intracellular end of TM6 triggered by G protein-binding highlight the essential role of G protein-binding in receptor activation. Nevertheless, all these simulations suggest that although the I92^3.40^N mutant does not induce a full active state for A_2A_AR, it can preserve the intermediate state that driven by the agonist through a hydrophilic interaction network; on the contrary, the WT receptor would return back to the inactive state within a short timescale once the agonist removed.

**Figure 4.**
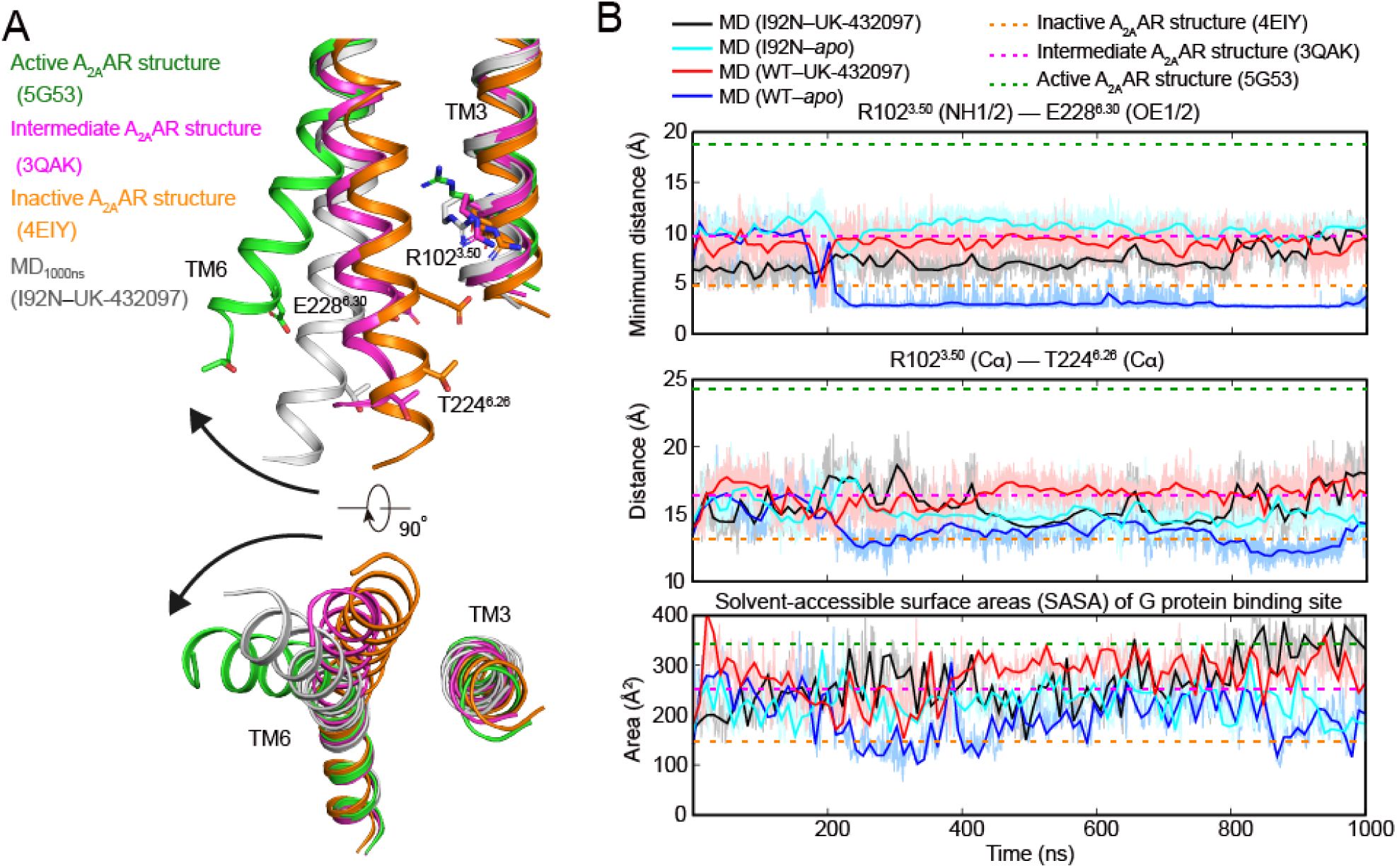
Conformational dynamics of the intercellular part of transmembrane helix 6 (TM6) during MD simulation. (**A**), Structural comparison of the intracellular half of TM6 between MD final snapshot and the released structure of A_2A_AR in different states. Inactive, intermediate, and active conformations are colored in orange, magenta and green, respectively (Carpenter et al., 2016; Liu et al., 2012; F. Xu et al., 2011). (**B**),Movements of TM6 during MD simulations: top, minimum distance between the charged non-hydrogen atoms of R102^3.50^ and E228^6.30^; middle, the Cα distance between R102^3.50^ and the intracellular tip of TM6 (T224^6.26^); bottom; the solvent-accessible surface area of G protein-binding sites, which consists of R102^3.50^, A105^3.53^, I106^3.54^, I200^5.61^, A203^5.64^, S234^6.36^ and L235^6.37^. The interface areas were calculated by FreeSASA (Mitternacht, 2016). Dashed horizontal lines indicate values for the released structure of A_2A_AR in different states (inactive state, orange; intermediate state, magenta; active, green).

## Discussion

In this study we determined the A_2A_AR constitutive active mutant I92^3.40^N in complex with agonist UK-432097 to a resolution of 3.8 Å. We identified that the mutation I92^3.40^N stabilizes a hydrophilic interaction network that preserved an intermediated state in the presence or removal of the agonist through MD simulations, whereas the WT receptor trends to move back to the inactive state without the presence of agonist. Our results indicated that both WT and mutant receptors can hardly equilibrate to the fully active conformation during the simulation in 1μs timescale, with or without agonist. This observation is consistent with the critical role of G protein binding in receptor activation, as revealed in previous structural and dynamic studies (Carpenter et al., 2016; Eddy et al., 2018; Ye et al., 2016). Alternatively, long-timescale accelerated/enhanced MD simulations have been developed to escape local energy minima and efficiently sample the full energy landscape (McRobb, Negri, Beuming, & Sherman, 2016). Additionally, there is another possibility that the current conformation observed in I92^3.40^N mutant structure is potentially affected by the lattice packing since a BRIL protein was inserted in the ICL3. Two BRIL proteins packing against each other in the lattice and a distortion is identified in the middle of each 4^th^ helix within the packing interface (Figure 2—figure supplement 1A-B). Such a distortion is not seen in previous BRIL structures that fused to ICL3 of GPCRs (Figure 2—figure supplement 2C-D), and it is possible that the lattice packing suppressed the position of BRIL and that pressure is immediately transferred to the TM6 of A_2A_AR through the rigid connection between receptor and fusion protein.

The residues involved in the common activation pathway are partially conserved within class A GPCRs, *e*.*g*., the W^6.48^ is located in a highly conserved CWxP motif, while the opposing position 3.40 is not very conserved but typically adopts a residue with short side-chain to fit the highly condensed region in the center region. Remarkably, several mutations on position 3.40 have been linked to dysfunctions or diseases, *i*.*e*., V509^3.40^A of thyrotropin receptor caused hyperthyroidism non-autoimmune (Duprez et al., 1994), I137^3.40^T of melanocortin receptor 4 caused obesity (Gu et al., 1999; Xiang et al., 2006), S127^3.40^F of vasopressin V2 receptor caused nephrogenic diabetes insipidus (Erdelyi et al., 2015), and L125^3.40^R of rhodopsin caused retinitis pigmentosa 4 (Dryja, 1992). These mutations may unbalance the activity of each receptor through either impulse the conformational transition (active) or disconnect the transition linkage (inactive). Our study laid the basis for understanding the mechanism for these disease-related mutations and can be effectively applied to future modeling studies for pharmacological or pathological purposes.

In summary, our research together with previous studies indicated the critical role of the transmission switch, and either agonist binding or specific mutations in the activation pathway may trigger receptor conformational change to achieve or maintain active or active-like state. Our research provides a general template to understand the mutation triggered conformational change and signal transduction though the combination of structural and computational biology, and highlights the single point mutation strategy may provide another routine to initiate signal transduction beside the classical agonist binding.

## Materials and Methods

### A_2A_AR construct design, expression and purification

Human wild-type A_2A_AR gene has 412 residues. The crystallization construct has replaced the ICL3 loop (residue K209-G218) by BRIL (thermostabilized apocytochrome b_562_ from *E. coli*) and cut off the C-terminal after A316 which hindering the protein crystallization. The modified A_2A_AR gene was cloned to pFastBac 1 vector containing HA signal peptide, FLAG epitope tag, and 3C protease cleavage site at the N-terminus and 10×His-tag at the C-terminus. Three mutations I92^3×40^N, L95^3×43^A and I238^6×40^Y were induced individually by overlap PCR to form constitutively active mutations. Recombinant baculoviruses expressing A_2A_AR WT or mutants were prepared using Bac-to-Bac system (Invitrogen). *Spodoptera frugiperda* 9 (*Sf9*) insect cells were grown in ESF921 medium, when *Sf9* cells density at 2∼3×10^6^ cells/ml can be infected by 1% (v/v) baculoviruses and harvested at 48 h after infection. The 1 L cells were collected by centrifugation, flash-frozen in liquid nitrogen, and stored at ™80 °C until further use. After 2 washes of hypotonic buffer (10 mM HEPES pH7.5, 10 mM MgCl_2_, 20mM KCl with EDTA-free protease-inhibitor cocktail tablets) and 3 washes of high salt buffer (10 mM HEPES pH7.5, 10 mM MgCl_2_, 20 mM KCl, 1 M NaCl with EDTA-free protease-inhibitor cocktail tablets), the cell pellets were collected and pre-treated with 4 mM theophylline (Sigma), 2.0 mg/ml iodoacetamide (Sigma), and EDTA-free protease-inhibitor cocktail tablets. After incubation for 30 min the cell membranes were solubilized by incubation in the presence of 50 mM HEPES, 500 mM NaCl, 1% N-Dodecyl-β-D-maltoside (DDM, Anatrace), 0.2% cholesterol hemisuccinate (CHS, Sigma), for 3h at 4 °C, The insoluble material was removed by centrifugation at 150,000 ×g and the supernatant was added to 1 ml pure TALON resin (Clontech) and 20 mM imidazole and rock slowly overnight at 4 °C. The resin was washed with 4×10 column volumes (CV) of wash buffer (25 mM HEPES pH7.5, 500 mM NaCl, 5% glycerol, 0.05% DDM, 0.01% CHS, 30 mM imidazole and 20 μM UK-432097), and eluted with 3 ml elution buffer (25mM HEPES pH7.5, 500 mM NaCl, 5% glycerol, 0.025% DDM, 0.005% CHS, 300 mM imidazole and 100 μM UK-432097). The elution was concentrated with 100 kDa molecular weight cut-off (MWCO) Amicon centrifugal ultrafiltration unit (Millipore).

### Thermal shift assay

CPM (N-([4-(7-diethylamino-4-methyl-3-coumarinyl)phenyl]maleimide) dye was dissolved in DMSO at 4 mg/mL as stock solution and diluted 20 times in CPM buffer (25 mM HEPES, pH 7.5, 500 mM NaCl, 5% (v/v) glycerol, 0.01% (w/v) DDM, 0.002% (w/v) CHS) before use. 1 μL of diluted CPM was added to the same buffer with approximately 0.5-2 μg receptor in a final volume of 50 μL. For receptors prepared for thermal shift assay, no compound was added during purification and each compound was only added in each CPM buffer to final concentration of 50 μM. The thermal shift assay was performed in a Rotor-gene real-time PCR cycler (Qiagen). The excitation wavelength was 365 nm and the emission wavelength was 460 nm. All assays were performed over a temperature range from 25°C to 85°C. The stability data were processed with GraphPad Prism.

### Crystallization

Purified A_2A_AR protein was cocrystallized with UK-432097 using lipid cubic phase (LCP) technology. The concentrated A_2A_AR was mixed the lipid [10% (w/w) cholesterol, 90% (w/w) monoolein] using a 1:1.5 (v/v) protein:lipid ratio to generate LCP mixture, then loaded 50 nl each well on 96 well plate and overlaid with 800 nl of different precipitant solution. LCP plates were stored at room temperature (18-20 °C). Diffracting quality crystals were grown in the condition of 100 mM Tris pH 8.2, 30% PEG400 and 0.4 M (NH_4_)_2_SO_4_. A_2A_AR™UK-432097 crystals were harvested using mesh grid loops (MiTeGen) and stored in liquid nitrogen before use.

### Data collection and model building

X-ray diffraction data were collected at the Japan synchrotron radiation SPring-8 on beaming line 45XU with automatic data collection program. Diffraction data were then collected with the 10 μm beam with 0.2 second exposures with an oscillation of 0.2° per frame. X-ray diffraction data were automatically processed with program KAMO (Yamashita, Hirata, & Yamamoto, 2018) and indexed and scaled using XDS (Kabsch, 2010). The structure was solved by molecular replacement with PHASER by using the solved A_2A_AR structures (PDB ID: 3QAK) as search models. Resulting model refinement and rebuilding were performed using Phenix (Adams et al., 2010) and Coot (Emsley, Lohkamp, Scott, & Cowtan, 2010). Statistics are provided in Table 1 and the final 3D pictures are prepared with PyMOL (The PyMOL Molecular Graphics System, Version 2.0 Schrödinger, LLC.).

### Molecular dynamic simulations

Molecular dynamic simulations were performed by Gromacs 2020.1 (Abraham et al., 2015). The WT A_2A_AR (UK-432097-bound A_2A_AR, PDB ID: 3QAK) and I92N mutant (crystal structure determined herein) were prepared and capped by the Protein Preparation Wizard (Schrodinger 2019-2). Two residues D52^2.50^ and D101^3.49^ were deprotonated, while other titratable residues were left in their dominant state at pH 7.0. The *apo* receptor or its complex with UK-432097 were embedded in a bilayer composed of 201 POPC lipids and solvated with 0.15 M NaCl in explicitly TIP3P waters using CHARMM-GUI Membrane Builder (Wu et al., 2014). The CHARMM36-CAMP force filed (Guvench et al., 2011) was adopted for protein, lipids and salt ions. The parameter of UK-432097 was generated using the CHARMM General Force Field (CGenFF) (Vanommeslaeghe et al., 2010) program version 2.4.0. The Particle Mesh Ewald (PME) (Darden, York, & Pedersen, 1993) method was applied with a cutoff of 10 Å and the bonds involving hydrogen atoms were constrained using LINCS algorithm (Hess, 2008). The MD simulation system was relaxed using the steepest descent energy minimization, followed by slow heating of the system to 310 K with restraints. The restraints were reduced gradually over 20 ns, with a simulation step of 1 fs. Finally, 1000 ns production run without restraints were carried out, with a time step of 2 fs in the NPT ensemble at 310 K and 1 bar using the v-rescale thermostat (Bussi, Donadio, & Parrinello, 2007) and the semi-isotropic Parrinello-Rahman barostat (Aoki & Yonezawa, 1992), respectively. The “gmx hbond” function within Gromacs was used to analyze hydrogen bond occupancies (applied criteria of donor-acceptor distance: 3.5 Å and 40°angle). The interface areas were calculated by FreeSASA(Mitternacht, 2016) using the Sharke-Rupley algorithm with a probe radius of 1.2 Å.

## Acknowledgments

This work was supported by the National Key Research and Development Program of China (2018YFA0507000, 2018YFA0507001), and the National Nature Science Foundation of China grant 31770898 (G.S.), 21704064 (Q.Z.), 31971178 (S.Z.) and the start-up funding by Fudan University (Q.Z.). We thank Prof. Fei Xu for helpful discussions.

## Author Contributions

M.C. made A_2A_AR mutations, expressed and purified proteins, crystallized and determined the structure; Q.Z. designed mutations, performed MD simulations, analyzed data and edited manuscript; Y.W. assisted insect cell culture; D.Y. assisted crystal data collection and process; S.Z. oversaw the project and edited manuscript; G.S. oversaw the crystallization and edited manuscript.

## Data availability

Atomic coordinates and structure factors for the A_2A_AR variant structure I92^3.40^N–UK-432097 has been deposited in the Protein Data Bank with identification code 7EZC. All data generated during this study are included in the manuscript and supporting files. Source data files have been provided for Figure 1 and Figure 1—figure supplement 1. Correspondence and requests for materials may be addressed to (Q.Z.) or (G.S.)

## Figures

**Figure 1—figure supplement 1.**
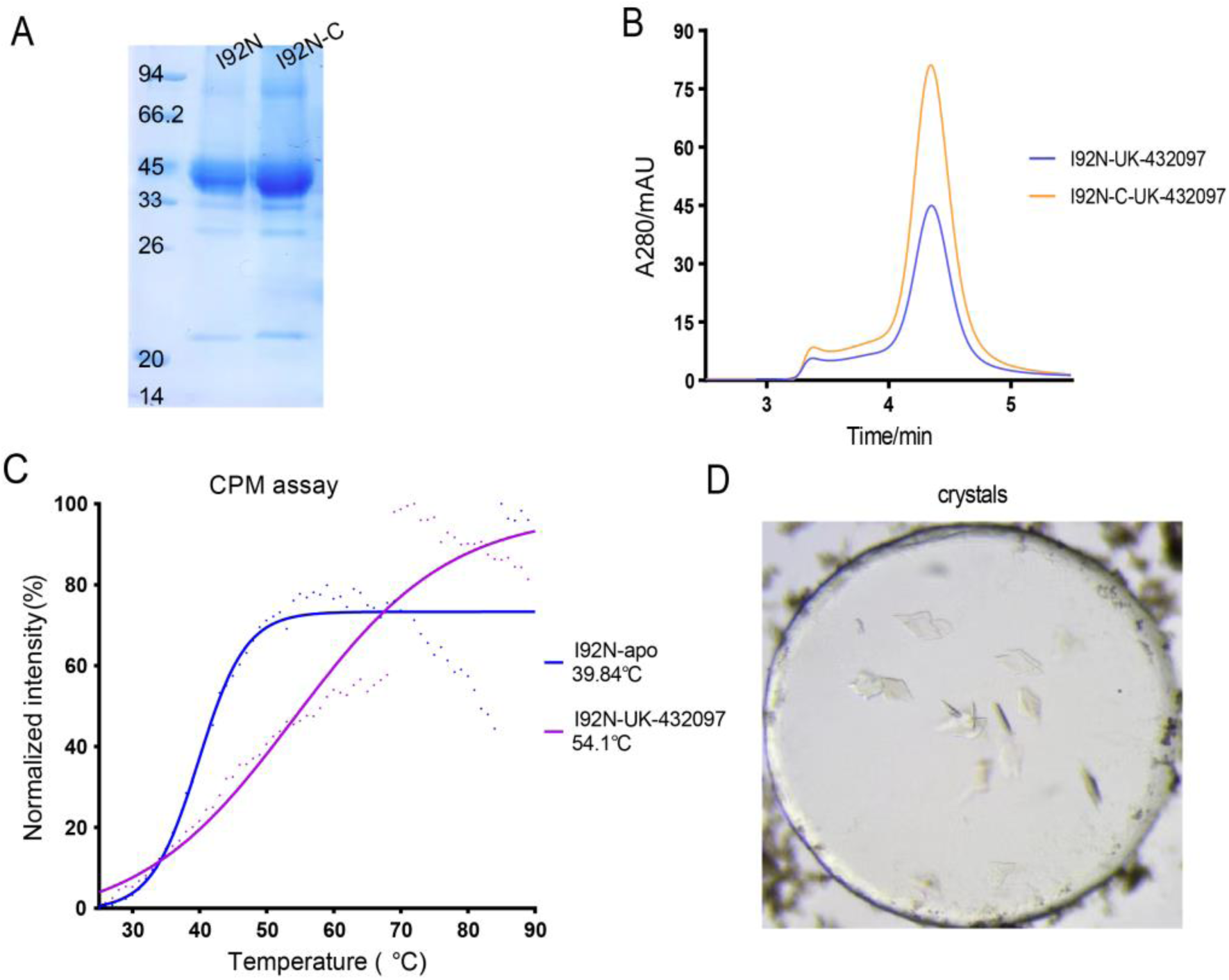
Purification and crystallization of I92N mutant with UK-432097. (**A**-**B**), SDS-PAGE and SEC profiles of purified mutant (I92N) and concentrated (I92N-C) I92N–UK-432097 complex. (**C**), Thermal-shift assay of mutant receptor in *apo* or in complex with UK-432097. Tm values are shown on the right. (**D**), Crystals of I92N–UK-432097 complex from LCP environment. **Source data 1**. Raw SDS-PAGE of A_2A_AR purified mutant (I92N) and concentrated (I92N-C) I92N–UK-432097 complex. **Source data 2**. Raw size-exclusion chromatography data of A_2A_AR purified mutant (I92N) and concentrated (I92N-C) I92N–UK-432097 complex. **Source data 3**. Raw CPM-based thermal-shift assay data of A_2A_AR I92N mutant receptor in *apo* or in complex with UK-432097.

**Figure 2—figure supplement 2.**
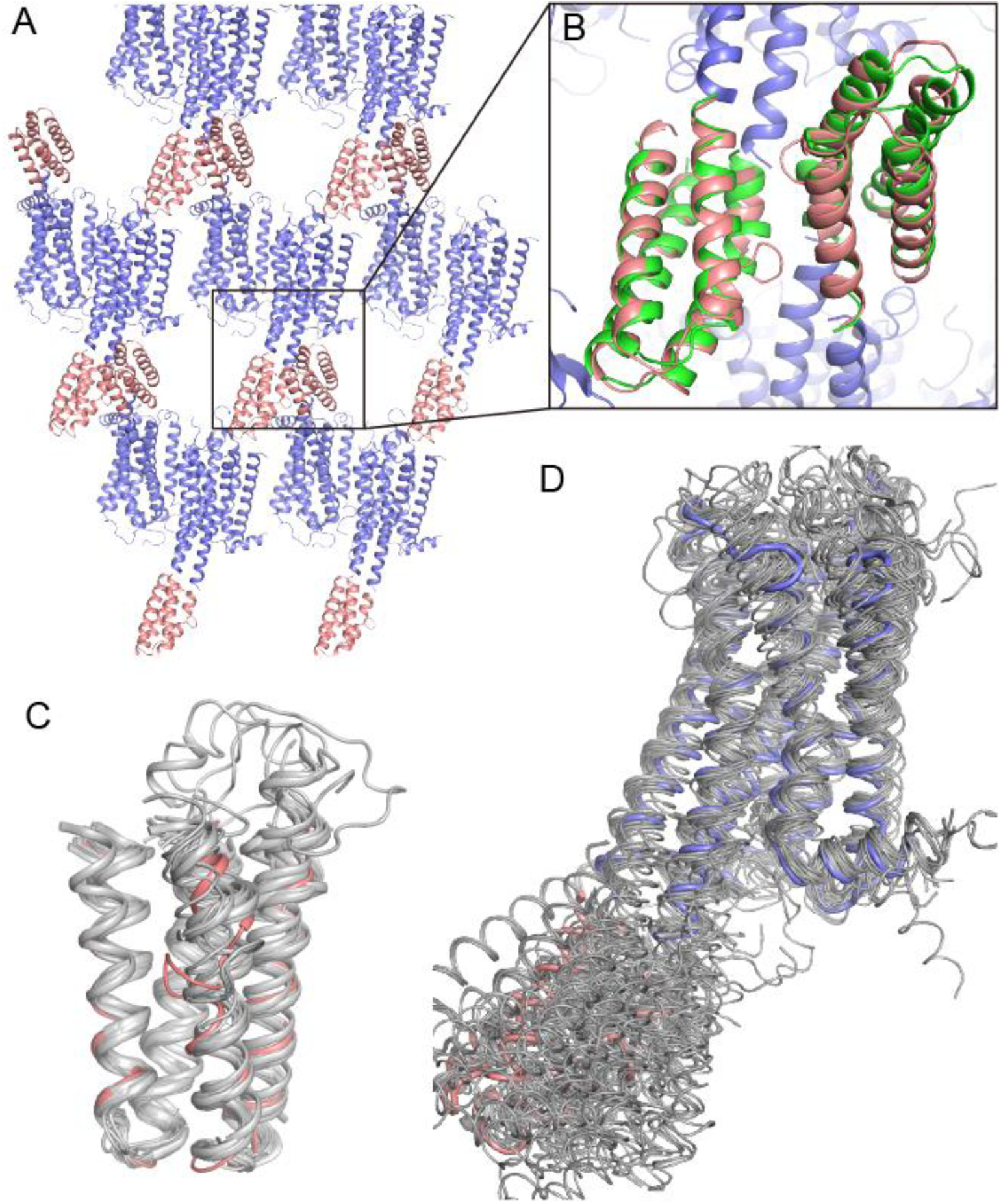
Crystal packing of I92N–UK-432097. (**A**-**B**), The lattice packing and BRIL-BRIL interface within the lattice. (**C**), Superposition of the BRIL proteins that used as fusion partner at the ICL3 of GPCRs. (**D**), Superposition on receptors of the GPCR-BRIL^ICL3^ fusion protein structures showed variant orientations for the ICL3-BRIL. PDBs used for alignments are PDBs: 4EIY, 4IAR, 4IB4, 4NTJ, 4PXZ, 4IAQ, 4Z35, 4Z34, 4Z36, 5UIG, 5IU4, 5IUA, 5N2R, 5MZJ, 5JTB, 5UEN, 6DRX, 6DRY, 6DRZ, 6RZ4, 6RZ6, 6RZ7, 6RZ8, 6RZ9, 6BQH, 6BQG, 6AQF, 6ZDR, 6ZDV and current structure. All these previous structures of GPCR-BRIL^ICL3^ fusion proteins showed varied orientations between the receptor and BRIL, and only the 5HT_2C_ receptor in complex with agonist is crystallized in a full active conformation.

## References

Abraham, M. J., Murtola, T., Schulz, R., Páll, S., Smith, J. C., Hess, B., & Lindahl, E. (2015). GROMACS: High performance molecular simulations through multi-level parallelism from laptops to supercomputers. SoftwareX, 1-2, 19–25. doi:10.1016/j.softx.2015.06.001

Adams, P. D., Afonine, P. V., Bunkoczi, G., Chen, V. B., Davis, I. W., Echols, N., Zwart, P. H. (2010). PHENIX: a comprehensive Python-based system for macromolecular structure solution. Acta Crystallogr D Biol Crystallogr, 66(Pt 2), 213–221. doi:10.1107/S0907444909052925

Aoki, K. M., & Yonezawa, F. (1992). Constant-pressure molecular-dynamics simulations of the crystal-smectic transition in systems of soft parallel spherocylinders. Physical Review A, 46(10), 6541–6549. doi:10.1103/physreva.46.6541

Ballesteros, J. A., & Weinstein, H. (1995). [19] Integrated methods for the construction of three-dimensional models and computational probing of structure-function relations in G protein-coupled receptors. In Receptor Molecular Biology (pp. 366–428).

Bussi, G., Donadio, D., & Parrinello, M. (2007). Canonical sampling through velocity rescaling. J Chem Phys, 126(1), 014101. doi:10.1063/1.2408420

Carpenter, B., Nehme, R., Warne, T., Leslie, A. G., & Tate, C. G. (2016). Structure of the adenosine A(2A) receptor bound to an engineered G protein. Nature, 536(7614), 104–107. doi:10.1038/nature18966

Darden, T., York, D., & Pedersen, L. (1993). Particle mesh Ewald: AnN·log(N) method for Ewald sums in large systems. J Chem Phys, 98(12), 10089–10092. doi:10.1063/1.464397

Dryja, T. P. (1992). Doyne Lecture. Rhodopsin and autosomal dominant retinitis pigmentosa. Eye (Lond), 6 (Pt 1), 1–10. doi:10.1038/eye.1992.2

Duprez, L., Parma, J., Van Sande, J., Allgeier, A., Leclere, J., Schvartz, C., et al. (1994). Germline mutations in the thyrotropin receptor gene cause non-autoimmune autosomal dominant hyperthyroidism. Nat Genet, 7(3), 396–401. doi:10.1038/ng0794-396

Eddy, M. T., Lee, M. Y., Gao, Z. G., White, K. L., Didenko, T., Horst, R., Wuthrich, K. (2018). Allosteric Coupling of Drug Binding and Intracellular Signaling in the A2A Adenosine Receptor. Cell, 172(1-2), 68–80 e12. doi:10.1016/j.cell.2017.12.004

Emsley, P., Lohkamp, B., Scott, W. G., & Cowtan, K. (2010). Features and development of Coot. Acta Crystallogr D Biol Crystallogr, 66(Pt 4), 486–501. doi:10.1107/S0907444910007493

Erdelyi, L. S., Mann, W. A., Morris-Rosendahl, D. J., Gross, U., Nagel, M., Varnai, P., … Hunyady, L. (2015). Mutation in the V2 vasopressin receptor gene, AVPR2, causes nephrogenic syndrome of inappropriate diuresis. Kidney International, 88(5), 1070–1078. doi:10.1038/ki.2015.181

Erlandson, S. C., McMahon, C., & Kruse, A. C. (2018). Structural Basis for G Protein-Coupled Receptor Signaling. Annu Rev Biophys, 47, 1–18. doi:10.1146/annurev-biophys-070317-032931

Garcia-Nafria, J., Lee, Y., Bai, X., Carpenter, B., & Tate, C. G. (2018). Cryo-EM structure of the adenosine A2A receptor coupled to an engineered heterotrimeric G protein. Elife, 7, 19. doi:10.7554/eLife.35946

Gu, W., Tu, Z., Kleyn, P. W., Kissebah, A., Duprat, L., Lee, J., Allison, D. B. (1999). Identification and functional analysis of novel human melanocortin-4 receptor variants. Diabetes, 48(3), 635–639. doi:10.2337/diabetes.48.3.635

Guvench, O., Mallajosyula, S. S., Raman, E. P., Hatcher, E., Vanommeslaeghe, K., Foster, T. J., Mackerell, A. D., Jr. (2011). CHARMM additive all-atom force field for carbohydrate derivatives and its utility in polysaccharide and carbohydrate-protein modeling. J Chem Theory Comput, 7(10), 3162–3180. doi:10.1021/ct200328p

Hess, B. (2008). P-LINCS: A Parallel Linear Constraint Solver for Molecular Simulation. J Chem Theory Comput, 4(1), 116–122. doi:10.1021/ct700200b

Kabsch, W. (2010). Xds. Acta Crystallogr D Biol Crystallogr, 66(Pt 2), 125–132. doi:10.1107/S0907444909047337

Latorraca, N. R., Venkatakrishnan, A. J., & Dror, R. O. (2017). GPCR Dynamics: Structures in Motion. Chem Rev, 117(1), 139–155. doi:10.1021/acs.chemrev.6b00177

Lebon, G., Edwards, P. C., Leslie, A. G., & Tate, C. G. (2015). Molecular Determinants of CGS21680 Binding to the Human Adenosine A2A Receptor. Mol Pharmacol, 87(6), 907–915. doi:10.1124/mol.114.097360

Lebon, G., Warne, T., Edwards, P. C., Bennett, K., Langmead, C. J., Leslie, A. G., & Tate, C. G. (2011). Agonist-bound adenosine A2A receptor structures reveal common features of GPCR activation. Nature, 474(7352), 521–525. doi:10.1038/nature10136

Lee, S., Nivedha, A. K., Tate, C. G., & Vaidehi, N. (2019). Dynamic Role of the G Protein in Stabilizing the Active State of the Adenosine A2A Receptor. Structure, 27(4), 703–712 e703. doi:10.1016/j.str.2018.12.007

Liu, W., Chun, E., Thompson, A. A., Chubukov, P., Xu, F., Katritch, V., Stevens, R. C. (2012). Structural basis for allosteric regulation of GPCRs by sodium ions. Science, 337(6091), 232–236. doi:10.1126/science.1219218

Manglik, A., Kim, T. H., Masureel, M., Altenbach, C., Yang, Z., Hilger, D., Kobilka, B. K. (2015). Structural Insights into the Dynamic Process of beta2-Adrenergic Receptor Signaling. Cell, 161(5), 1101–1111. doi:10.1016/j.cell.2015.04.043

McRobb, F. M., Negri, A., Beuming, T., & Sherman, W. (2016). Molecular dynamics techniques for modeling G protein-coupled receptors. Curr Opin Pharmacol, 30, 69–75. doi:10.1016/j.coph.2016.07.001

Mitternacht, S. (2016). FreeSASA: An open source C library for solvent accessible surface area calculations. F1000Res, 5, 189. doi:10.12688/f1000research.7931.1

Nygaard, R., Zou, Y., Dror, R. O., Mildorf, T. J., Arlow, D. H., Manglik, A., Kobilka, B. K. (2013). The dynamic process of beta(2)-adrenergic receptor activation. Cell, 152(3), 532–542. doi:10.1016/j.cell.2013.01.008

Rasmussen, S. G., DeVree, B. T., Zou, Y., Kruse, A. C., Chung, K. Y., Kobilka, T. S., Kobilka, B. K. (2011). Crystal structure of the beta2 adrenergic receptor-Gs protein complex. Nature, 477(7366), 549–555. doi:10.1038/nature10361

Schonegge, A. M., Gallion, J., Picard, L. P., Wilkins, A. D., Le Gouill, C., Audet, M., Bouvier, M. (2017). Evolutionary action and structural basis of the allosteric switch controlling beta2AR functional selectivity. Nat Commun, 8(1), 2169. doi:10.1038/s41467-017-02257-x

Song, W., Yen, H. Y., Robinson, C. V., & Sansom, M. S. P. (2019). State-dependent Lipid Interactions with the A2a Receptor Revealed by MD Simulations Using In Vivo-Mimetic Membranes. Structure, 27(2), 392–403 e393. doi:10.1016/j.str.2018.10.024

Thal, D. M., Glukhova, A., Sexton, P. M., & Christopoulos, A. (2018). Structural insights into G-protein-coupled receptor allostery. Nature, 559(7712), 45–53. doi:10.1038/s41586-018-0259-z

Vanommeslaeghe, K., Hatcher, E., Acharya, C., Kundu, S., Zhong, S., Shim, J., Mackerell, A. D., Jr. (2010). CHARMM general force field: A force field for drug-like molecules compatible with the CHARMM all-atom additive biological force fields. J Comput Chem, 31(4), 671–690. doi:10.1002/jcc.21367

Venkatakrishnan, A. J., Deupi, X., Lebon, G., Tate, C. G., Schertler, G. F., & Babu, M. M. (2013). Molecular signatures of G-protein-coupled receptors. Nature, 494(7436), 185–194. doi:10.1038/nature11896

Wacker, D., Wang, S., McCorvy, J. D., Betz, R. M., Venkatakrishnan, A. J., Levit, A., Roth, B. L. (2017). Crystal Structure of an LSD-Bound Human Serotonin Receptor. Cell, 168(3), 377–389 e312. doi:10.1016/j.cell.2016.12.033

Weis, W. I., & Kobilka, B. K. (2018). The Molecular Basis of G Protein-Coupled Receptor Activation. Annu Rev Biochem, 87, 897–919. doi:10.1146/annurev-biochem-060614-033910

White, K. L., Eddy, M. T., Gao, Z. G., Han, G. W., Lian, T., Deary, A., Stevens, R. C. (2018). Structural Connection between Activation Microswitch and Allosteric Sodium Site in GPCR Signaling. Structure, 26(2), 259–269 e255. doi:10.1016/j.str.2017.12.013

Wu, E. L., Cheng, X., Jo, S., Rui, H., Song, K. C., Davila-Contreras, E. M., Im, W. (2014). CHARMM-GUI Membrane Builder toward realistic biological membrane simulations. J Comput Chem, 35(27), 1997–2004. doi:10.1002/jcc.23702

Xiang, Z., Litherland, S. A., Sorensen, N. B., Proneth, B., Wood, M. S., Shaw, A. M., Haskell-Luevano, C. (2006). Pharmacological characterization of 40 human melanocortin-4 receptor polymorphisms with the endogenous proopiomelanocortin-derived agonists and the agouti-related protein (AGRP) antagonist. Biochemistry, 45(23), 7277–7288. doi:10.1021/bi0600300

Xu, F., Wu, H., Katritch, V., Han, G. W., Jacobson, K. A., Gao, Z. G., Stevens, R. C. (2011). Structure of an agonist-bound human A2A adenosine receptor. Science, 332(6027), 322–327. doi:10.1126/science.1202793

Xu, Y., Wang, Y., Wang, Y., Liu, K., Peng, Y., Yao, D., Song, G. (2019). Mutagenesis facilitated crystallization of GLP-1R. Iucrj, 6(Pt 6), 996–1006. doi:10.1107/S2052252519013496

Yamashita, K., Hirata, K., & Yamamoto, M. (2018). KAMO: towards automated data processing for microcrystals. Acta Crystallogr D Struct Biol, 74(Pt 5), 441–449. doi:10.1107/S2059798318004576

Yang, D., Zhou, Q., Labroska, V., Qin, S., Darbalaei, S., Wu, Y., Wang, M. W. (2021). G protein-coupled receptors: structure-and function-based drug discovery. Signal Transduct Target Ther, 6(1), 7. doi:10.1038/s41392-020-00435-w

Ye, L., Van Eps, N., Zimmer, M., Ernst, O. P., & Prosser, R. S. (2016). Activation of the A2A adenosine G-protein-coupled receptor by conformational selection. Nature, 533(7602), 265–268. doi:10.1038/nature17668

Zhou, Q., Yang, D., Wu, M., Guo, Y., Guo, W., Zhong, L., Zhao, S. (2019). Common activation mechanism of class A GPCRs. Elife, 8, e50279. doi:10.7554/eLife.50279

